## Supplementary Information for "More than the sum of its parts: investigating episodic memory as a multidimensional cognitive process"

### Supplementary Methods

##### Visual perception task

To behaviorally quantify each participant's perception, the accuracy of the responses was used. On average, we ended up with 205 trials associated with high perception (range of 106 to 255, SD = 33) and 64 trials associated with low perception (range of 27 to 119, SD = 20).

##### Sustained attention task

To behaviorally quantify each participant's sustained attention, in addition to the response accuracy, the standard deviation of response time for correct responses divided by the mean response time and the accuracy of the responses were used. We divided the standard deviation response time by the mean response time to control for variability for participants' processing speed. On average, we ended up with 91 trials associated with high sustained attention and 91 trials associated with low sustained attention (range of 44 to 96, SD = 8 for both conditions).

##### Selective attention task

To behaviorally quantify each individual's selective attention, in addition to the response accuracy, the difference of standard deviation response time for the invalid and in the valid conditions will be divided by the mean response time during both conditions. We divided the difference by the mean response time to control for variability for participants' processing speed and to highlight the impact of invalidity on performance. On average, we ended up with 114 trials associated with high selective attention and 114 trials associated with low selective attention (range of 84 to 119, SD = 6 for both conditions).

#### Episodic memory task

To behaviorally quantify each participant's item memory, item memory d' was used. Furthermore, to behaviorally quantify each participant's attended context memory (regardless of whether it was the color or the scene), attended context memory d' was computed as Z (proportion of "match" responses to contexts that matched those presented at encoding) – Z (proportion of match responses to contexts that mismatched those shown at encoding). On average, we ended up with 160 item hits (range of 76 to 216, SD = 30) and 56 item misses (range of 21 to 113, SD = 23). On average, we ended up with 104 context correct decisions (range of 59 to 177, SD = 30) and 55 context incorrect decisions (range of 16 to 116, SD = 30).

#### The control analyses for validating the transfer learning results

The first control analysis tested the importance of using *meaningful* sources predicted to support memory encoding as opposed to using *any* type of source to improve memory decoding performance. In this regard, we used randomly generated noise as an external source and investigated how much leveraging information from that external source could improve memory prediction performance compared to investigating memory as a unidimensional process. As stated above, from each source, we extracted CSP-based features from the voltage and the power of theta, alpha, beta, and gamma frequency bands and then used the RCSP algorithm to leverage the source information on the encoding data. We repeated the same process here, but the source data (from which we extracted the CSP-based features) was randomly generated noise. In more detail, for every participant, we computed random values within the range of values identified in our EEG dataset (across all source tasks for that participant) separately for voltage and the power of each selected frequency band. We generated 500 events using this approach with 250 of them randomly labeled as “low” and the other 250 labeled as “high”. We then performed the transfer learning procedure (described above) to transfer this source to the target.

The second control analysis tested the importance of using the training portion of the encoding data to make the necessary adjustments when transferring a source to the target. As previously mentioned, $\alpha$ and $\beta$ are the regularized parameters that need to be calculated (i.e., during cross validation) to effectively transfer the source information to the target data by making the appropriate adjustment to the selected CSP-based features. Previously, we mentioned how choosing $\alpha = 0$ and $\beta = 0$ would result in the typical unidimensional approach for the classification of the target task which totally ignores the information from the source. Similarly, by choosing $\alpha = 1$ and $\beta = 0$, transfer learning completely ignores the target information and makes no adjustments to transfer the source information to the target data. And that’s what we did in this control analysis to inspect the importance of using the training portion of the encoding data to make the appropriate adjustments to effectively transfer a source to the target.

**Confirming dissociability of the 3 sources**

We combined the events of all these three tasks and performed a 3-class classification. The label of each event was the task it belonged to (regardless of whether that event was a high or low performance event). We used a one versus rest voting approach to generalize the binary classification into a 3-class classification[4]. To elaborate, the first binary classifier would classify sustained attention events vs other events (perception and selective attention collapsed as a single class). Similarly, the second binary classifier would classify selective attention events vs other events, and the third binary classifier would classify perception events vs other events. The classifier evidence scores for these three binary classifiers would be combined to collectively predict which task an event belonged to. For each binary classification analyses performed here, we used the same procedure we used for other analyses which we described above (i.e., extracting CSP-based features from voltage and the power of different frequency bands, using a combination of the filter and wrapper methods to select the best 5 features among the 40 filtered features prior to training a naïve Bayes classifier).

#### Supplementary Notes

##### Behavioral Results

Behavioral performance associated with each task is shown in Supplementary Table 1.

##### The classification results to predict attended context memory success

We found that transfer learning significantly enhanced our ability to predict attended context memory success across participants compared to the traditional approach/unidimensional approach (from 68.5% to 78.3%) [$t(42) = 11.220, p < 0.001, \text{one} - \text{tailed}, d = 0.56$; Supplementary Fig. 8]. In addition, we inspected how much memory classification performance would improve by adding each source in a stepwise manner rather than including all 3 sources simultaneously. We added the sources in all six possible orders and the patterns of results were similar and thus, we report the average findings. We found that there was a 5.2% performance improvement after the first source was added, regardless of order. There was a 2.9% performance improvement once the second source was added followed by a 1.7% performance improvement once the third source was added (Supplementary Fig. 8). Statistically, we found that adding each source significantly improved memory classification performance [$all\ ts > 2.785, all\ ps < 0.005, \text{one} - \text{tailed}, all\ ds > 0.10$]. Moreover, the extent to which classification performance increased by adding a source decreased with each step [step 2 improvement compared to step 1: $t(42) = 1.946, p = 0.029, \text{one} - \text{tailed}, d = 0.50$ and step 3 improvement compared to step 2: $t(42) = 1.210, p = 0.117, \text{one} - \text{tailed}, d = 0.31$]. It is also worth mentioning that all the events included in attended context memory classification were item hits. As a result, it is not surprising that attended context memory classification accuracy is not as high as item memory classification accuracy.

**Investigating selective attention as a multidimensional process**

In these analyses, we used selective attention as the target domain and used visual perception, sustained attention, and episodic encoding as the sources. We found that investigating selective attention as a multidimensional process using transfer learning significantly enhanced our ability to predict the selective attention level across participants compared to the traditional approach/unidimensional approach (from 66.8% to 76.4%) [$t(42) = 13.351, p < 0.001, \text{one} - \text{tailed}, d = 1.97$; Supplementary Fig. 9]. In addition, we inspected how much classifying the selective attention brain states would improve by adding each source in a stepwise manner rather than including all 3 sources simultaneously. We found that adding the first and second source significantly improved the classification performance, regardless of the order in which the sources were added [$all\ ts > 4.588, all\ ps < 0.001, \text{one} - \text{tailed}, all\ ds > 0.41$]. However, unlike the memory classification results, the order in which the sources were added mattered for how much the third source impacts the selective attention classification performance. Specifically, when added as the third source, while visual perception [$t(42) = 4.049, p < 0.001, \text{one} - \text{tailed}, d = 0.32$] and sustained attention [$t(42) = 4.329, p < 0.001, \text{one} - \text{tailed}, d = 0.37$] significantly improved the selective attention classification performance, adding episodic encoding as the third source did not lead to a significant improvement [$t(42) = 1.170, p = 0.124, \text{one} - \text{tailed}, d = 0.08$]. This suggests that episodic encoding cannot explain unique variance of selective attention-related neural activity while the information related to visual perception and sustained attention has already been leveraged. Furthermore, we investigated whether there would be a diminishing return every step a new source is added. Again, the order in which the sources were added mattered. We first compared the improvements at the first and second steps. We found that when visual perception or sustained attention was added as the first source, the extent to which classification performance increased by adding a source significantly decreased [$all\ ts > 3.099, all\ ps < 0.002, \text{one} - \text{tailed}, all\ ds > 0.70$]. However, when episodic encoding was added as the first source, the extent to which classification performance increased by adding a source did not significantly decrease [$all\ ts < 1.673, all\ ps > 0.050, \text{one} - \text{tailed}, all\ ds < 0.40$]. We then compared the improvements at the second and third steps. We found that when visual perception or sustained attention was added as the second source, the extent to which classification performance increased by adding a source significantly decreased [$all\ ts > 3.800, all\ ps < 0.001, \text{one} - \text{tailed}, all\ ds > 0.88$]. However, when episodic encoding was added as the second source, the extent to which classification performance increased by adding a source did not significantly decrease [$all\ ts < 1.532, all\ ps > 0.065, \text{one} - \text{tailed}, all\ ds < 0.37$].

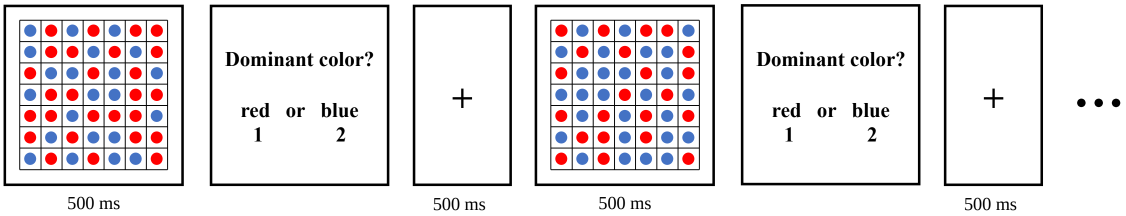


**Supplementary Fig. 1. The perception task.**

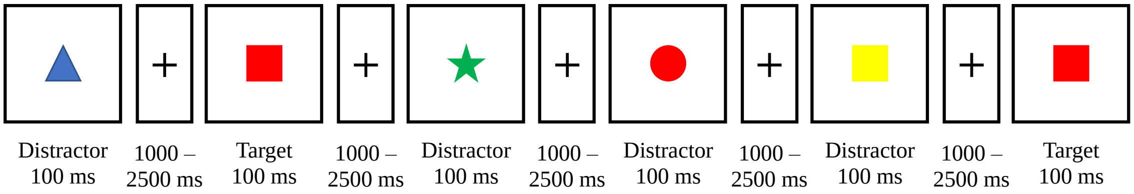


**Supplementary Fig. 2. The Conjunctive Continuous Performance Test-Visual (CCPT-V) task.** This task was used to assess the participants' level of sustained attention.

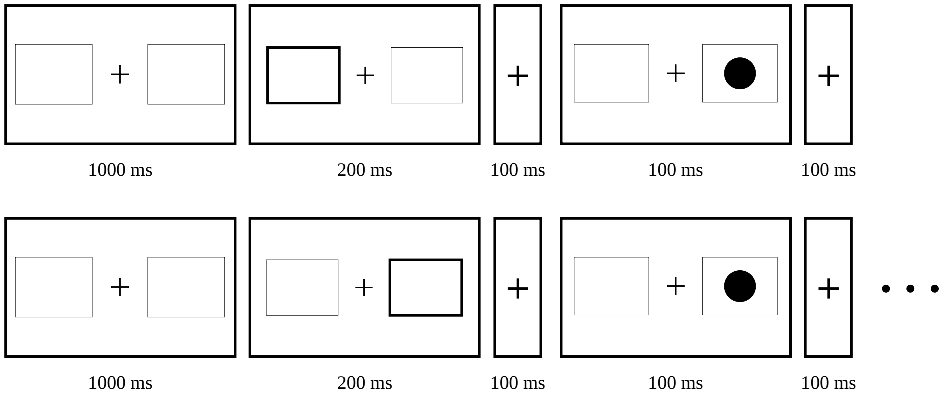


**Supplementary Fig. 3. The Spatial Cued-Identification Task (SCIT) task.** This task was used to assess the participants' level of selective attention.

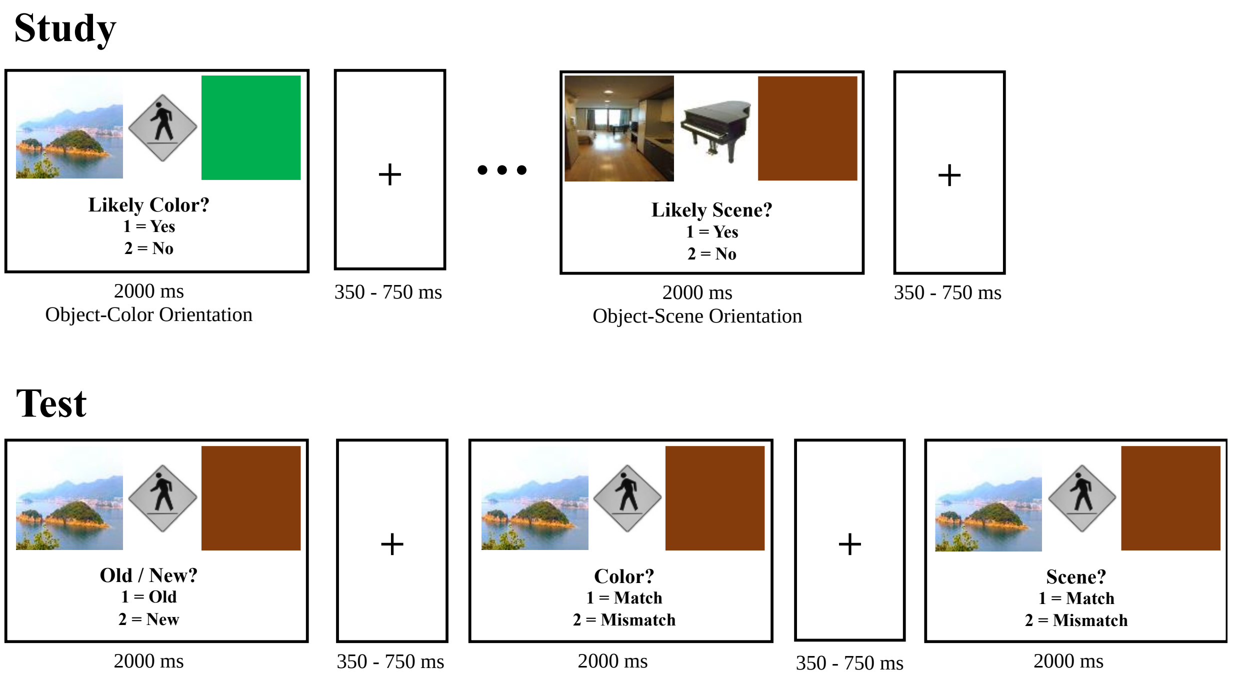


**Supplementary Fig. 4. Task design for the episodic memory study.** The scenes in this figure are taken from Creative Commons.

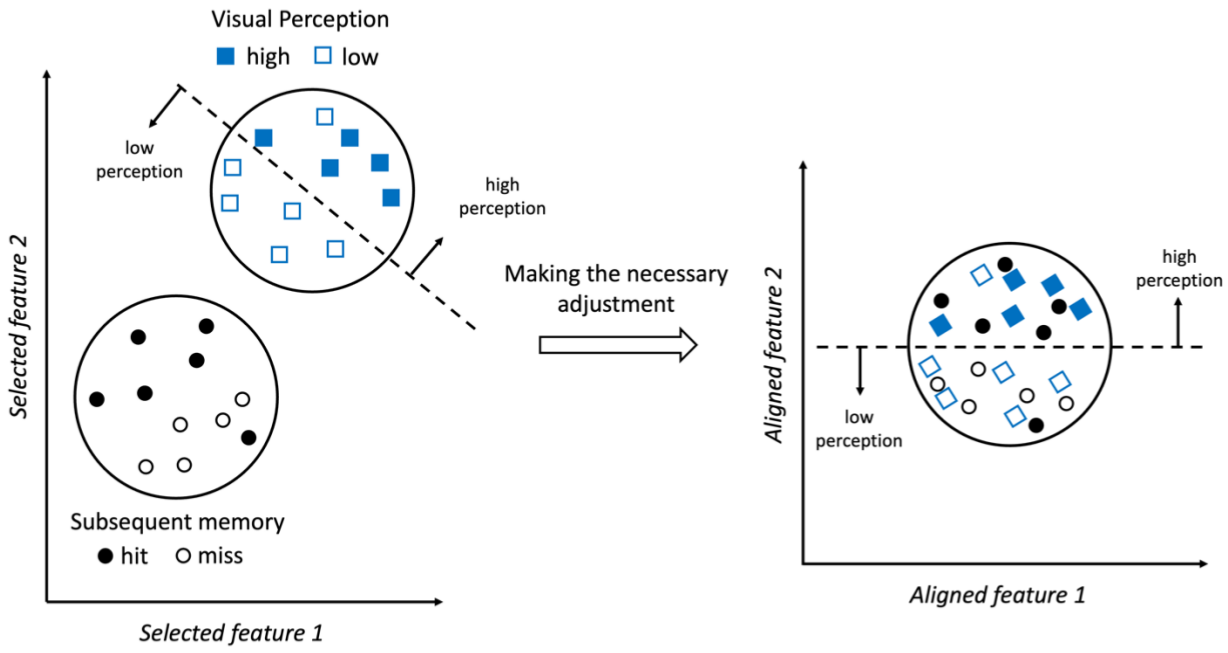


**Supplementary Fig. 5. An illustration of how transfer learning makes the necessary adjustments to transfer a source to the target.** The example in this figure is just for demonstrative purposes to provide an intuition of how transfer learning works. Specifically, when training a high vs. low perception performance classifier, we realize which CSP-based features (i.e., selected features 1 and 2 in this figure) can best distinguish high vs low perception events. The perception events are the blue squares and are shown in the selected 2-dimensional feature space. The decision boundary to separate high and low perception brain states is shown with a dashed line. Next, the same selected features will be extracted from the training portion of the encoding data and the encoding events will be projected into the selected feature space. However, when trying to determine the encoding events' perception level, the current decision boundary would label all the encoding events as low perception, suggesting the need to use transfer learning to make necessary adjustments. As such, transfer learning uses the information from the perception and the encoding data to project the data from both domains into a new *aligned* feature space that can effectively predict the perception level during encoding events.

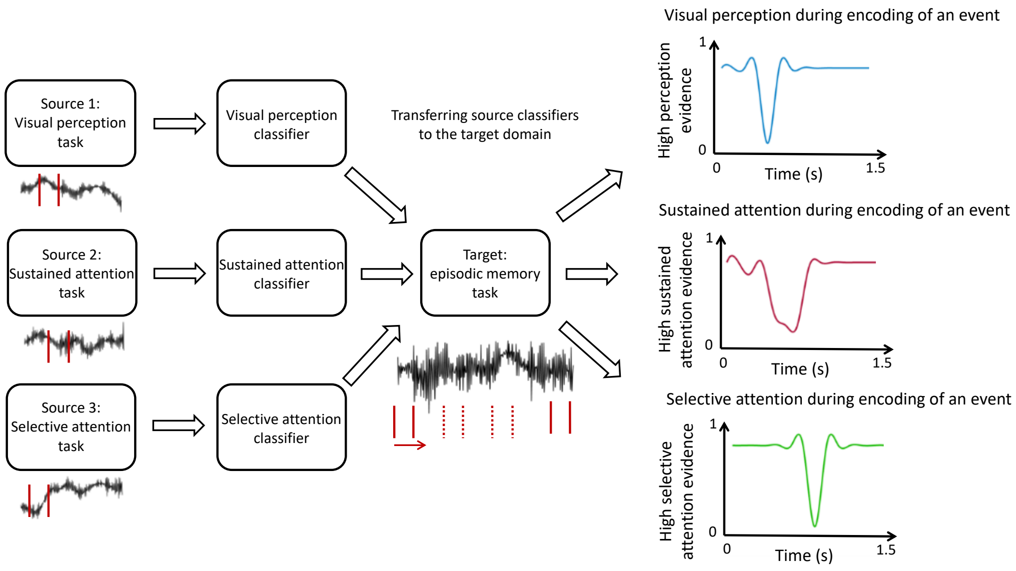


**Supplementary Fig. 6. An illustration of obtaining a temporal map of how high the brain state associated with each source is during an encoding event.** The high vs. low performance classifier for each source was trained using an optimal 200 ms time window across participants. These optimal 200 ms time windows could be different across the three sources but for each source, the same 200 ms time window was used across all participants. Using a sliding time window approach, the information regarding the high vs low levels of each source was transferred to different encoding periods for each event. This allowed us to determine the level of each source during different encoding periods for each event. Notably, the evidence for the low brain state for each source would be 1 - the evidence for the high brain state for that source. For example, if there is 0.8 evidence for high perception for the first 200 ms during an event's episodic encoding, there will be 0.2 evidence for low perception, but we have not showed that here to keep the figure as simple as possible.

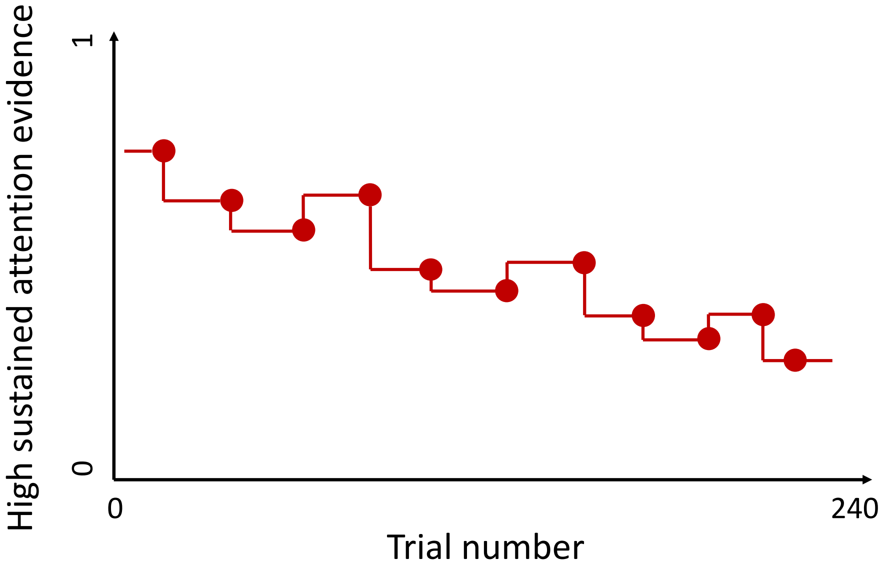


**Supplementary Fig. 7.** The interpolation approach to associate neural evidence to all 240 trials for each memory condition (i.e., hits and misses). In this figure, the red circles are the trials that are associated with the misses for that participant.

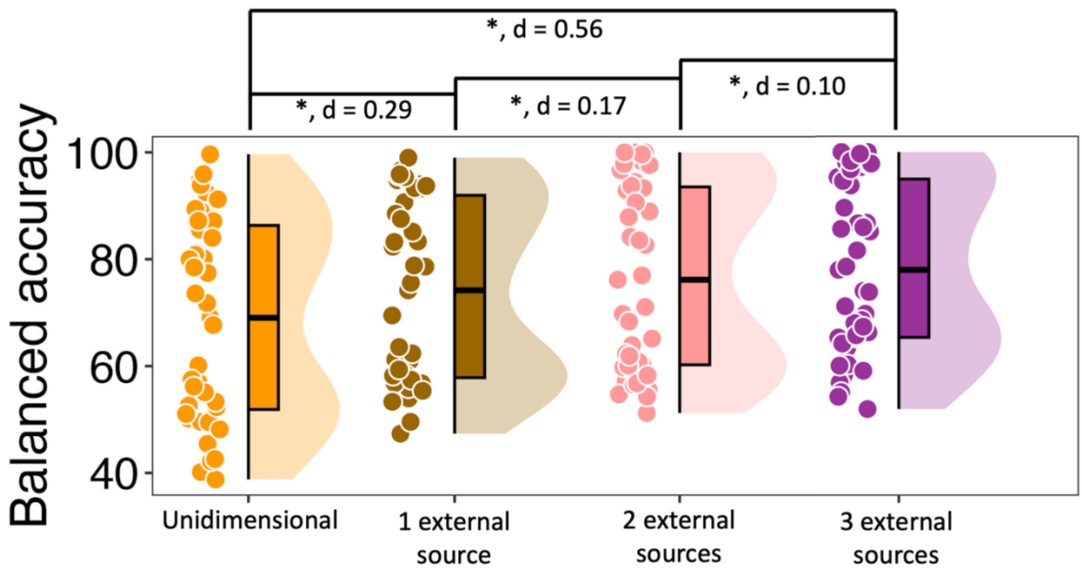


**Supplementary Fig. 8. Attended context memory classification results.** Comparisons of balanced accuracy for classifying attended context memory brain states as a function of how many of the sources are included (averaged across all six possible orders) during classification. The asterisk signs show significant differences between the conditions and the associated Cohen's d is shown for each comparison.

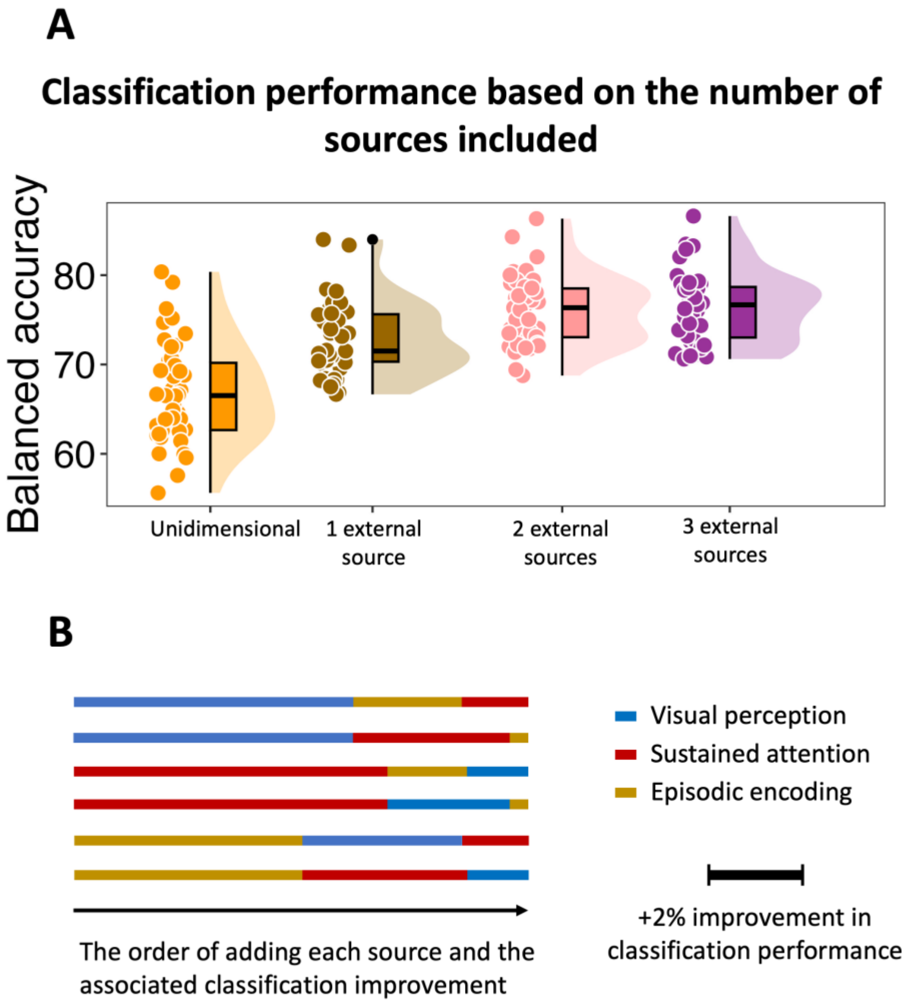


**Supplementary Fig. 9. Investigating selective attention as a multidimensional process. (A)** Comparisons of balanced accuracy for classifying selective attention brain states as a function of how many of the sources are included during classification. In the analysis associated with this figure, visual perception was added first, followed by sustained attention and then episodic encoding. **(B)** The extent to which each added source improved the classification performance depending on the order in which the sources were added. Note that for memory classification results, we did not show this pattern as the results were very similar across the six possible orders the sources could be added. However, given the nature of the sources and the target in this particular analysis, the order in which the sources were added mattered and thus, the associated result for each possible order is shown. The length of each line represents the extent to which adding that source improved the classification performance.

| | |
|---|---|
| Item d' | $2.08 \pm 0.063\ (SE)$ |
| Attended context d' | $1.65 \pm 0.064\ (SE)$ |
| Perception | $74.64\% \pm 1.27\%\ (SE)$ |
| Sustained attention mean response time | $451.0\ ms \pm 5.2\ ms\ (SE)$ |
| Sustained attention performance | $92.29\% \pm 0.93\%\ (SE)$ |
| Selective attention mean response time valid | $338.6\ ms \pm 6.9\ ms\ (SE)$ |
| Selective attention mean response time invalid | $386.9\ ms \pm 6.5\ ms\ (SE)$ |
| Selective attention performance | $90.70\% \pm 1.28\%\ (SE)$ |

**Supplementary Table 1. The behavioral performance on the tasks associated with the sources and the target.** SE stands for standard error.
